# Optimizing Language Model Embeddings to Voxel Activity Improves Brain Activity Predictions

**DOI:** 10.1101/2025.09.18.676935

**Authors:** Anuja Negi, Christine Tseng, Anwar O Nunez-Elizalde, Xue Lily Gong, Fatma Deniz

## Abstract

Recent studies have shown that contextual semantic embeddings from language models can accurately predict human brain activity during language processing. However, most studies use contextual embeddings with the same context length and model layer for all voxels, potentially overlooking meaningful variations across the brain. In this study, we investigate whether optimizing contextual embeddings for individual voxels improves their ability to predict brain activity during reading. We optimize embeddings for each voxel by selecting the best-predicting context length, model layer, or both. We perform this optimization with two different types of stimuli (isolated sentences and narratives), and quantify the performance gains of optimized embeddings over standard fixed embeddings. Our results show that voxel-specific optimization substantially improves the prediction accuracy of contextual semantic embeddings. These findings demonstrate that voxel-specific contextual tuning provides a more accurate and nuanced account of how the contextual semantic information is represented across the cortex.

## Introduction

Recent research has demonstrated that contextual semantic embeddings extracted from text-based transformer language models can be used to accurately predict human brain activity during language processing (Toneva and Wehbe 2019; Anderson et al. 2021; Caucheteux and King 2022; Caucheteux, Gramfort, and King 2022; Goldstein et al. 2022; Karamolegkou, Abdou, and Søgaard 2023; Yamashita, Kubo, and Nishimoto 2025). These models of brain activity, also called brain encoding models, commonly use contextual semantic embeddings with a fixed context length and model layer that are chosen to maximize the average prediction accuracy across all voxels. However, this global optimization assumes that the same configuration is optimal for the entire brain, overlooking meaningful variations in how different regions process language. This limitation is significant, as different brain regions have been shown to represent language at distinct timescales(Lerner et al. 2011; Chen et al. 2024; Goldstein et al. 2025; Yamashita, Kubo, and Nishimoto 2025). Indeed, recent work has shown that contextual embeddings with different context lengths best predict different brain regions (Tikochinski et al. 2025). This prompts the question: *Does optimizing language model embeddings for individual voxels yield more accurate models of brain responses to language?*

In this study, we optimize contextual semantic embeddings for individual voxels and investigate if this improves brain encoding performance (i.e., the “prediction accuracy” or the “brain alignment score” of the encoding model). We perform this optimization for two commonly used types of stimuli, isolated sentences and narratives. We use functional magnetic resonance imaging (fMRI) data from participants reading text from both stimulus conditions. We first define a baseline contextual semantic embedding by selecting the context length and model layer that maximizes the mean contextual encoding performance across voxels. Next, we optimize contextual embeddings for each voxel in three ways: selecting the optimal context length, optimal model layer, or jointly optimizing both. Finally, we evaluate the benefits of voxel-specific optimization by quantifying the gains in brain encoding performance over the baseline contextual semantic embedding. All analyses are performed separately for each participant and stimulus condition.

Our results reveal two main findings. First, optimizing context length and model layer individually for each voxel significantly improved encoding performance in over 98% of voxels within the language network (Fedorenko et al. 2010). This result was consistent across both the isolated sentences and narratives conditions. Second, the largest improvements were consistently observed in bilateral angular gyrus, medial parietal cortex, and inferior frontal gyrus, all of which are involved in processing higher-level linguistic features (e.g., syntax, sentence- and paragraph-level semantics). Importantly, joint optimization of both context length and model layer yielded greater performance gains than optimizing either alone. These findings demonstrate the value of moving beyond a one-size-fits-all approach when using language model embeddings to predict brain responses. Our results contribute to ongoing efforts to align biological and artificial language representations, and underscore the need for fine-grained modeling to capture the complexity of language processing in the brain.

## Related Works

### Modeling language processing in the brain with language models

Our work builds on recent advances in using language models to predict human brain responses during language processing. Early studies established that static lexical embeddings can predict brain responses during both listening and reading (Huth et al. 2016; Deniz et al. 2019). More recent studies demonstrated that contextual embeddings from transformer-based language models can predict brain responses with even higher accuracy than lexical embeddings (Toneva and Wehbe 2019; Abdou et al. 2021; Antonello et al. 2021; Oota, Alexandre, and Hinaut 2022; Lamarre, Chen, and Deniz 2022). Additionally, encoding performance in language-selective brain regions improves as the size of the language model increases (Schrimpf et al. 2021; Caucheteux and King 2022). Recent studies also suggest that there is a representational alignment between transformer-based language models and language processing in the brain (Sucholutsky et al. 2023). Specifically, intermediate layers of models such as BERT and GPT-2 have been shown to align with specific stages of language processing in the brain, suggesting a layered correspondence between computational and cognitive hierarchies (Toneva and Wehbe 2019; Schrimpf et al. 2021). When using contextual embeddings to predict brain responses, prior studies either chose an arbitrary context length and model layer for all voxels (LeBel, Jain, and Huth 2021), or they optimized the context length and model layer by maximizing the average encoding performance across voxels (Toneva and Wehbe 2019; Anderson et al. 2021; Schrimpf et al. 2021; Caucheteux and King 2022; Caucheteux, Gramfort, and King 2022; Goldstein et al. 2022; Yamashita, Kubo, and Nishimoto 2025). However, results from neuroscience studies suggest that each voxel may have a different optimal context length and model layer (see next section). Our study addresses this gap by optimizing the context length and model layer specifically for each voxel.

### Language processing in the brain

Our work builds on neuroscience studies showing that different brain regions represent different types of linguistic information during language processing. First, neuroscience studies have shown that brain representations of lower-level linguistic features (e.g., lexical semantics) are spatially distinct from representations of higher-level linguistic features (e.g., paragraph-level semantics) during language comprehension (Lerner et al. 2011; Chang, Nastase, and Hasson 2022; Chen et al. 2024; Goldstein et al. 2025; Yamashita, Kubo, and Nishimoto 2025). Because lower-level linguistic features occur on the scale of individual words while higher-level linguistic features occur across multiple words, these results indicate that distinct brain regions represent information integrated across different amounts of context. This suggests that voxels in different brain regions may be optimally predicted by embeddings with different context lengths. Indeed, a recent study showed that embeddings with different context lengths significantly predict different sets of brain regions during language comprehension(Tikochinski et al. 2025). Second, language is hypothesized to be processed hierarchically in the brain, and neuroscience studies have identified distinct brain regions representing different levels of the language processing hierarchy (Price 2012; Fedorenko, Ivanova, and Regev 2024). Studies of language models suggest they are also organized hierarchically, with earlier layers representing lower-level linguistic features and later layers representing higher-level linguistic features (Tenney et al. 2019). Specific layers of LLMs have been shown to align to specific stages of language processing in the brain (Schrimpf et al. 2021), suggesting that voxels across different brain regions may be best predicted by embeddings from different model layers. Most prior studies using contextual embeddings have focused on either isolated sentences or narratives, which differ in the amount of context contained in the text. Recent work shows that context level affects both the signal-to-noise ratio (SNR) of recorded brain responses and linguistic representation (Deniz et al. 2023). This suggests that optimal context length and model layer may vary with the type of language stimuli used in the study. Our study addresses these gaps by optimizing the context length and model layer for individual voxels across isolated sentences and narratives.

## Methodology

### Brain Imaging Dataset

We used a subset of the data from Deniz et al. (2023) that we preprocessed with a custom preprocessing pipeline. Blood oxygen level-dependent (BOLD) responses were collected using fMRI from four right-handed, English-speaking participants (two males and two females) under four stimulus conditions with different amounts of linguistic context. We excluded data from one participant (male) due to low BOLD SNR and analyzed data from the other three participants. To align with prior brain encoding studies, we used data from two stimulus conditions: isolated sentences (medium context) and narratives (long context). We also included data from the isolated words (no context) condition as a control. Stimulus words were presented individually using Rapid Serial Visual Presentation (RSVP; (Forster 1970)), and different stimulus conditions were presented in different scanning runs (see Deniz et al. (2023) for details). Each condition had four unique runs (∼ 10 minutes each), one of which was repeated twice. The narratives condition consisted of four stories from The Moth Radio Hour (Huth et al. 2016) (“under-theinfluence,” “souls,” “life,” and “wheretheressmoke”). The isolated sentences condition comprised sentences randomly sampled from the 10 model estimation narratives used in Huth et al. (2016). The isolated words condition consisted of words randomly sampled without replacement from the same set of narratives. Of the four runs in each condition, three runs were used for training voxelwise encoding models (training set: 1149 TRs in narratives, 957 TRs in isolated sentences, and 930 TRs in isolated words; 1 TR=2.0045 sec; TR: repetition time), and the two repeated runs were averaged across repetitions and used for testing (test set: 311 TRs in narratives, 311 TRs in isolated sentences, and 310 TRs in isolated words; 1 TR=2.0045 sec).

More details about the dataset and preprocessing are provided in Appendix Section A.1.

### Feature Spaces

To model different types of information in the stimulus, we constructed feature spaces that captured low-level visual information, lexical semantic, and contextual semantic information.

#### Low-level visual feature space

To account for variance in the BOLD response due to low-level visual information during reading, motion energy features were calculated for each frame of the stimulus. The motion energy features reflect the spatiotemporal structure of the visual stimulus and are computed using a spatiotemporal Gabor pyramid (for details, see (Nishimoto et al. 2011)).

#### Lexical semantic features space

To capture context-independent semantic information, we extracted static lexical semantic features from the English1000 model Huth et al. (2016). Each word was represented as a 985-dimensional vector based on the word co-occurrence statistics in a large text corpus. This lexical feature space served as a baseline for evaluating the added value of the contextual embeddings.

#### Contextual semantic feature spaces

To capture contextual semantic information, we extracted contextual semantic features from two different transformer-based language models: English BERT (Devlin et al. 2019) and GPT-2 (Radford et al. 2019). English BERT is a bidirectional encoder with 12 transformer layers, each with a hidden size of 768. GPT-2 is a unidirectional decoder model with 12 layers and a hidden size of 768. For each model, we varied the input context length (number of preceding tokens) across six values: 0, 5, 10, 20, 40, and 80 tokens. For every combination of the model, context length, and model layer, we extracted contextual features from the hidden state activations. Each word was represented by the mean of all token-level embeddings corresponding to that word and its preceding context tokens (e.g., for a context length of 5 tokens, each word *w* was represented by the mean of all token-level embeddings of *w* and the 5 tokens preceding *w*).

### Optimizing Contextual Semantic Features for Each Voxel

To account for the fact that different voxels represent different linguistic features and timescales of information, we implemented voxel-specific optimization of the contextual semantic features. For each voxel, we defined the optimal contextual semantic embedding as the embedding that yielded the highest contextual semantic encoding performance. We performed this optimization under three settings: (1) selecting the optimal context length while fixing the model layer (optimal CL), (2) selecting the optimal model layer while fixing the context length (optimal layer), and (3) jointly optimizing both context length and model layer (optimal CL and layer).

The results reported in the main paper are based on contextual semantic features extracted from English BERT. The results and analyses using GPT-2 are included in Appendix Section C.

### Voxelwise Encoding Modeling (VEM)

To evaluate the relationship between contextual semantic features and stimulus context, we fitted multiple linearized encoding models for every voxel in each participant’s brain (Wu, David, and Gallant 2006; Naselaris et al. 2011). Each linearized voxelwise encoding model (VEM) consisted of a low-level visual feature space (motion energy) and a semantic feature space (either a lexical or contextual semantic feature space). A separate VEM was fitted for each voxel, participant, stimulus condition, lexical semantic feature, and contextual semantic feature (for each combination of context length and model layer). Before model fitting, all features and brain responses were z-scored across time. The features were then downsampled to match the fMRI acquisition rate using 3-lobe Lanczos interpolation. To account for the delayed hemodynamic response, we applied a finite impulse response (FIR) filter with four delays (2, 4, 6, and 8 seconds) to each feature space. For every encoding model, the two feature spaces (low-level visual and semantic) were fit as a single joint model to the training set BOLD responses using banded ridge regression (Nunez-Elizalde, Huth, and Gallant 2019; La Tour et al. 2022). Banded ridge regression is a special case of Tikhonov regression where each feature space is assigned a different regularization parameter. Fivefold cross-validation was used to optimize the regularization parameters. The prediction accuracy of the joint model was quantified as the Pearson correlation coefficient (*r*) between the predicted and recorded BOLD responses in the test set, and it is referred to as the joint model encoding performance in this paper. To assess the contribution of each feature space to the joint model encoding performance, we computed the “split” encoding performance for each feature space. The split encoding performance decomposes the joint model encoding performance into the unique contribution of each feature space by computing a variation of the product measure (La Tour et al. 2022). For concision, we refer to the split encoding performance of the lexical semantic feature space as the “lexical encoding performance” and the split encoding performance of the contextual semantic feature space as the “contextual encoding performance” in this paper.

All reported results are based on voxels within regions that are known to support language comprehension (Fedorenko et al. 2010), which we refer to here as language-selective regions. To define subject-specific ROIs, each participant’s cortical surface was transformed into the FsAverage6 template from Freesurfer (https://surfer.nmr.mgh.harvard.edu/) using surface-based registration. The group-level language mask (shown in Appendix Section A.2) was then projected onto individual brains to extract participant-specific language regions.

For reproducibility, please refer to Appendix Section D for implementation details.

## Results

### Contextual Encoding Performance Across Context Lengths, Model Layers, and Stimulus Conditions

To investigate the effect of optimizing the context length and model layer for every voxel, we first defined a baseline contextual semantic embedding against which to compare the optimized contextual semantic embeddings. To do this, we selected a fixed context length and model layer by optimizing the contextual encoding performance across all voxels in a set of language-selective brain regions (Fedorenko et al. 2010). We first fit a separate encoding model for every context length, model layer, voxel, participant, and stimulus condition. We then averaged the contextual encoding performance across all language-selective voxels in all participants. Finally, we defined the baseline contextual semantic embedding as the embedding with the context length and model layer that resulted in the highest average contextual encoding performance. Figure 2 shows the averaged contextual encoding performance for all combinations of context length, model layer, and stimulus condition.

**Figure 1.**
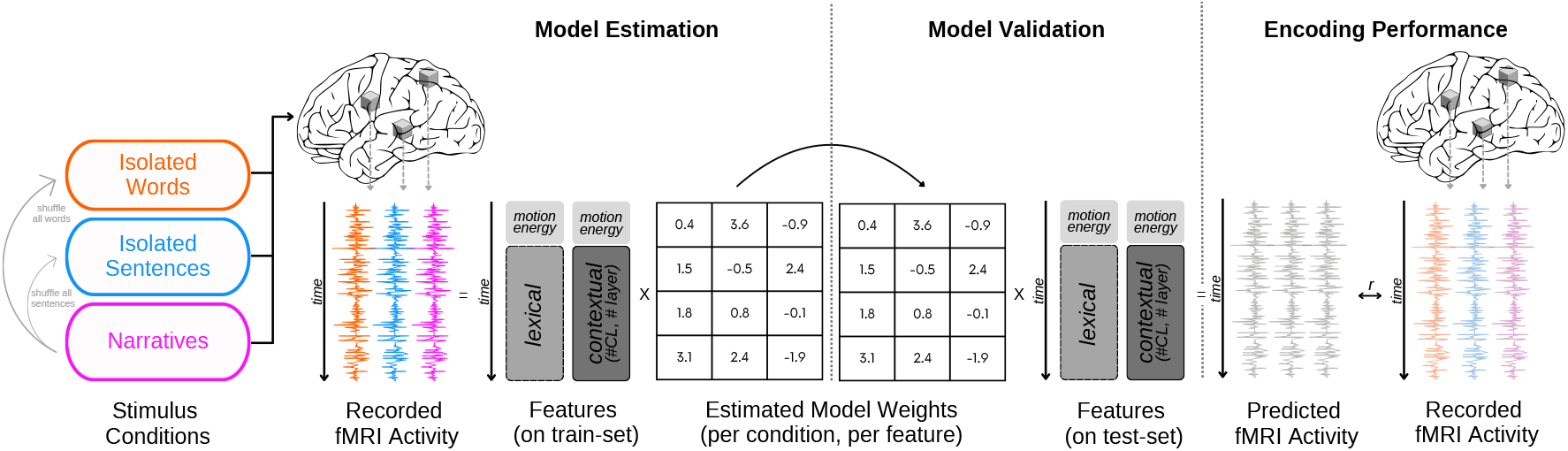
Voxelwise encoding modeling pipeline. Participants read language stimuli under three conditions: isolated words, isolated sentences, and narratives, while their fMRI responses were recorded. Each stimulus word was mapped onto multiple feature spaces: low-level visual features (using motion energy), lexical semantic features (word embeddings independent of context), and contextual semantic features derived from a pretrained language model. Contextual features were extracted from each model layer and computed for varying input context lengths (CL) of 0, 5, 10, 20, 40, 60, and 80 tokens. All features were aligned with the BOLD signal using a finite impulse response (FIR) model to account for hemodynamic delays. A joint encoding model combining semantic and low-level features was fitted separately for each voxel, subject, and stimulus condition using banded ridge regression. Encoding performance was calculated on held-out data by correlating the predicted and recorded fMRI activity.

**Figure 2.**
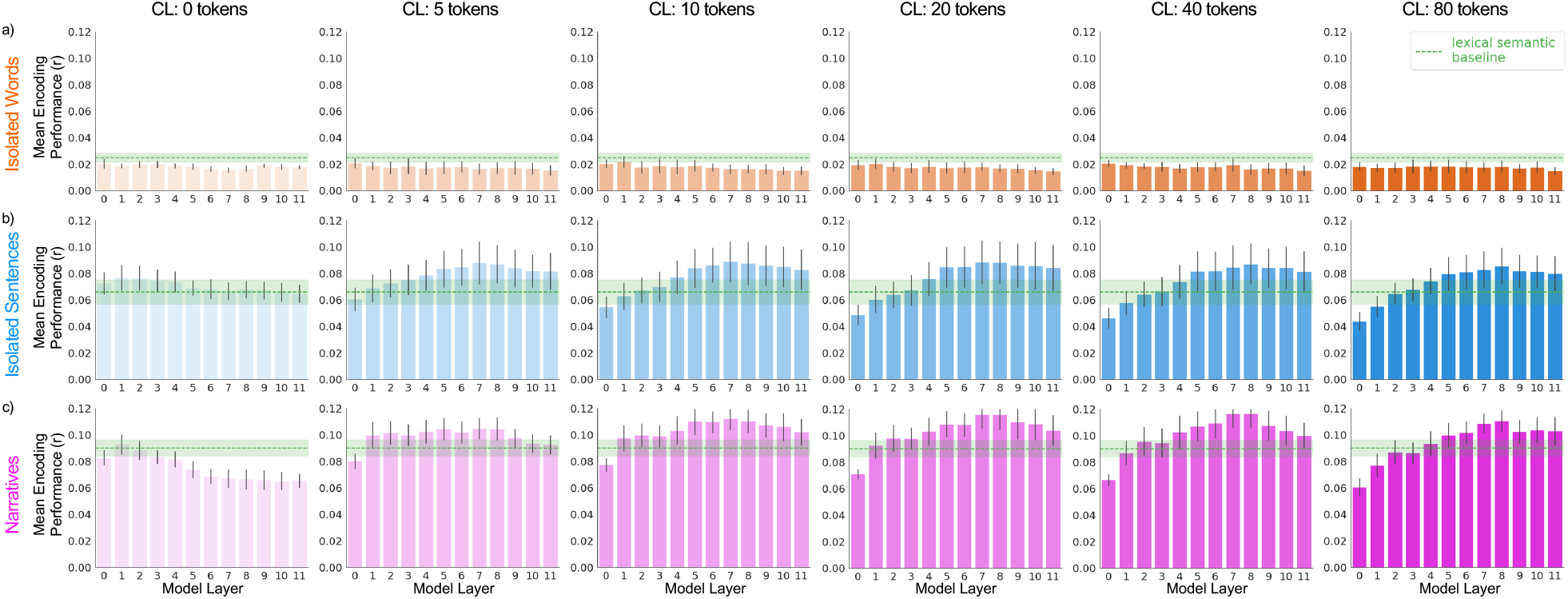
Brain encoding performance across context lengths, model layers, and stimulus conditions. We evaluated how context length and model layer influence brain encoding performance across three stimulus conditions. Voxelwise encoding models (VEMs; see Methods) were trained using a joint feature space that combined either lexical semantic features or contextual semantic features with low-level visual features (see Methods). For each model layer, contextual embeddings were computed using context lengths of 0, 5, 10, 20, 40, and 80 preceding tokens, as shown from left to right (columns 1–6). The rows correspond to the stimulus conditions: (a) isolated words (orange), (b) isolated sentences (blue), and (c) narratives (pink). Each subplot reports the mean encoding performance of the contextual semantic feature space (contextual encoding performance) averaged across language-selective regions (Fedorenko et al. 2010) and participants. Error bars indicate the standard error of the mean across participants. The dotted green line represents the encoding performance of the lexical semantic model (lexical encoding performance); the shaded green band indicates the standard error of the mean across participants. Overall, middle to late model layers (layers 5 to 9) and longer context lengths led to better encoding performance in the isolated sentences and narratives conditions compared to the lexical semantic model. In contrast, the single words condition shows no benefit from increased context length, with performance consistently below the lexical semantic model.

In the isolated words condition (Fig. 2a), contextual semantic embeddings performed worse than the lexical semantic embedding for all context lengths and model layers. In the isolated sentences condition (Fig. 2b), the average contextual encoding performance for context length 0 was comparable to the lexical encoding performance in all layers. For non-zero context lengths, contextual encoding performance surpassed the lexical encoding performance in the middle and late layers. Contextual encoding performance peaked around layers 7, 8, and 9 for all non-zero context lengths, creating a U-shaped performance curve across layers. The optimal context length and model layer were context length 20, model layer 8. In the narratives condition (Fig. 2c), the average contextual encoding performance for context length 0 was comparable to the lexical encoding performance in the early layers, but substantially worse than the lexical encoding performance in the middle and late layers. For non-zero context lengths, contextual encoding performance surpassed the lexical encoding performance in middle and late layers, with slightly larger improvement than in the isolated sentences condition (mean increase of 44.2% in narratives vs. 33.7% in isolated sentences). Similar to the isolated sentences condition, contextual encoding performance peaked around layers 7, 8, and 9, creating a U-shaped performance curve across layers. The optimal context length and model layer were context length 20, model layer 8.

The U-shaped curves observed in the isolated sentences and narratives conditions are consistent with the layer trends reported in previous studies (Toneva and Wehbe 2019; Caucheteux and King 2022; Schrimpf et al. 2021). Our results show that this trend also depends on the stimulus condition. In the isolated words condition, the stimuli consisted of individual words randomly sampled from a set of narratives. Because there was no coherent context, participants likely treated each word independently and ignored previously seen words. This explains the lower performance of contextual semantic embeddings (which contain meanings of previous words) compared to the lexical semantic embedding across all layers in this condition.

In the isolated sentences and narratives conditions, the presence of contextual information in the stimulus is reflected in the improvement of the contextual encoding performance over the lexical encoding performance. Interestingly, we see that the improvement in the narratives condition is slightly higher than the improvement in the isolated sentences condition. This may be because the narratives condition contains more long-range contextual information than the isolated sentences condition (sentence length median=13 words, min=5 words, max=40 words). Prior research has shown that different layers of BERT encode distinct types of linguistic information: early layers capture lower-level lexical features, whereas deeper layers represent higher-level linguistic structures such as co-reference and long-distance dependencies (Tenney, Das, and Pavlick 2019; Jawahar, Sagot, and Seddah 2019; Rogers, Kovaleva, and Rumshisky 2021). Thus, the deeper layers may be better matched to the statistics of the narratives condition.

Overall, we find that the optimal context length and model layer when considering all language-selective voxels is context length 20, layer 8 for both the isolated sentences and narratives conditions. We use this as the baseline contextual semantic embedding in the following analyses. In addition, we find that contextual encoding performance varies with context length, model layer, and stimulus condition, reflecting interactions between the linguistic context in the stimulus and the representational depth of the language model.

Appendix Section C shows the consistency of these results with another language model with a different architecture (GPT-2).

### Optimizing Contextual Features for Each Voxel Improves Brain Encoding Performance

To see whether optimizing the context length and model layer for every voxel improves contextual encoding performance, we compared the contextual encoding performance for optimized contextual semantic embeddings to that for the baseline contextual semantic embedding (context length 20, model layer 8). For each voxel, we optimized either the context length, model layer, or both by taking the context length and/or model layer that maximized contextual embedding performance for that voxel. Figure 3 shows all three comparisons (Optimal context length, Optimal model layer, Optimal context length and layer) for voxels in language-selective regions for both the isolated sentences and narratives conditions.

**Figure 3.**
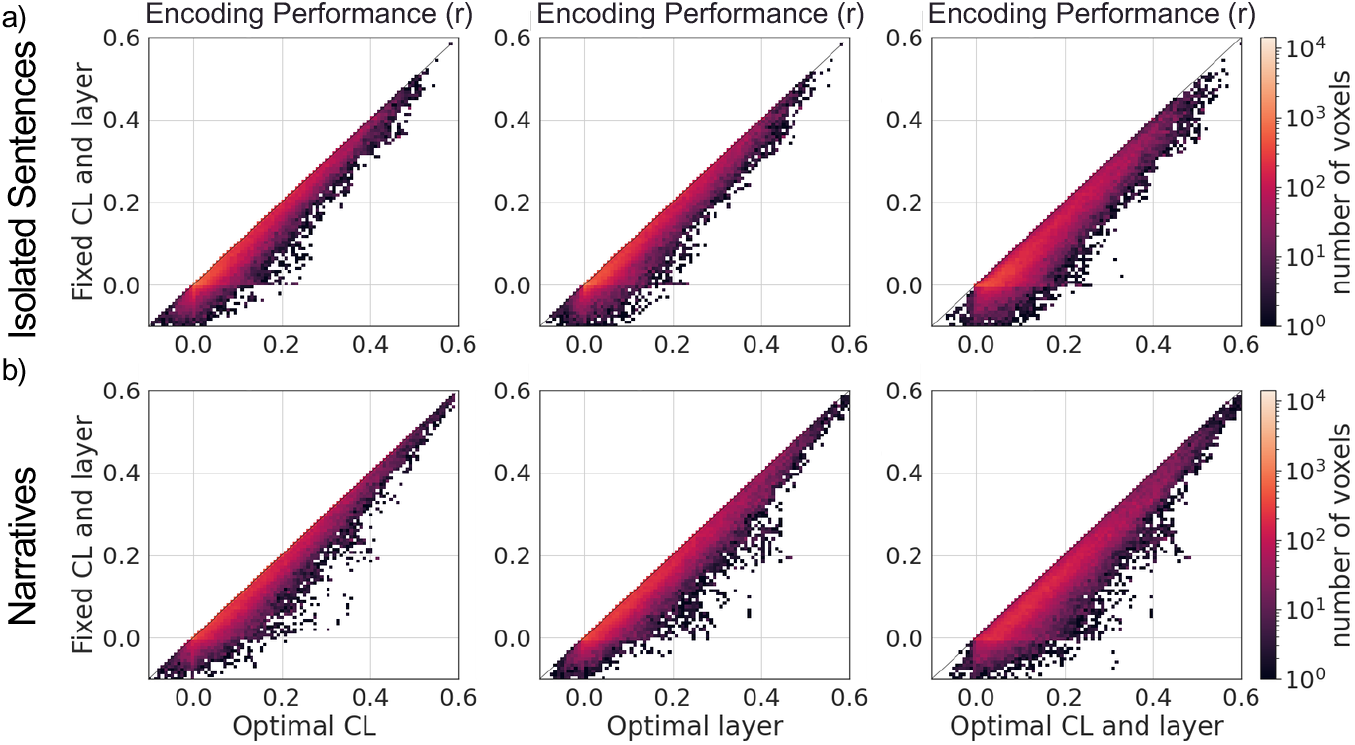
Gains in encoding performance from contextual feature optimization. To evaluate whether tailoring contextual features to each voxel improves encoding performance, we compared voxelwise encoding model (VEM; see Methods) performance using contextual embeddings with fixed context length and layer versus embeddings optimized per voxel. Specifically, we tested three settings: (1) using the optimal context length for each voxel while keeping layer fixed (column 1), (2) using the optimal layer with a fixed context length (column 2), and (3) jointly optimizing both context length and layer per voxel (column 3). Results are shown for two stimulus conditions: (a) isolated sentences and (b) narratives. Each subplot is a 2D histogram where the color indicates the number of voxels (colorbar) that exhibit a given pair of encoding performances (x: optimized features, y: fixed features). Voxels below the diagonal line show improved performance with voxel-specific optimization. Across both conditions and all settings, we observe a substantial number of voxels benefiting from optimized contextual representations. This demonstrates that optimizing context length and model layer to individual voxels leads to more accurate brain predictions. All results in this figure use contextual embeddings from the English BERT model, with fixed features defined as a context length of 20 and layer 8. The results shown are for Participant 1. The results for the other participants and the GPT-2 model are provided in Appendix Sections B and C, respectively.

#### Optimal context length

We created contextual embeddings with optimal context lengths by keeping the model layer fixed at 8 and selecting the context length that gave the highest contextual encoding performance for each voxel. We then compared contextual encoding performance between this optimized context length embedding and the baseline contextual embedding (context length 20, model layer 8). We observed improvements in ∼ 85% of language-selective voxels in the isolated sentences condition (mean improvement=0.02±0.02, max improvement=0.24) and ∼ 80% of language-selective voxels in the narratives condition (mean improvement=0.03±0.03, max improvement=0.29).

#### Optimal model layer

We created contextual embeddings with optimal model layers by keeping the context length fixed at 20 and selecting the model layer that gave the highest contextual encoding performance for each voxel. We then compared contextual encoding performance between this optimized model layer embedding and the baseline contextual embedding (context length 20, model layer 8). We observed improvements in ∼ 88% of language-selective voxels in the isolated sentences condition (mean improvement=0.03±0.03, max improvement=0.23) and ∼ 89% of language-selective voxels in the narratives condition (mean improvement=0.02±0.03, max improvement=0.31).

#### Optimal context length and layer

We constructed contextual embeddings with optimal context lengths and model layers by jointly selecting the context length and model layer that gave the highest contextual encoding performance for each voxel. We then compared contextual encoding performance between this optimized contextual embedding and the baseline contextual embedding (context length 20, model layer 8). We observed improvements in approximately 98% of language-selective voxels in the isolated sentences condition (mean improvement=0.05±0.03, max improvement=0.3) and 98% of language-selective voxels in the narratives condition (mean improvement=0.05±0.04, max improvement=0.38).

These results clearly demonstrate that optimizing contextual embeddings for individual voxels substantially increases contextual encoding performance.

Results for other participants and for GPT-2 model are provided in Appendix Sections B and C.

### Improved Encoding Performance Across Language Regions

To evaluate how much voxel-specific optimization of contextual embeddings improves contextual encoding performance across different brain regions, we plotted the improvement in contextual encoding performance between optimized and fixed contextual semantic embeddings across the cortex. Figure 4 shows this improvement for every voxel in the language-selective regions on a flattened FsAverage6 cortical surface for one participant (see Appendix section B for other participants).

**Figure 4.**
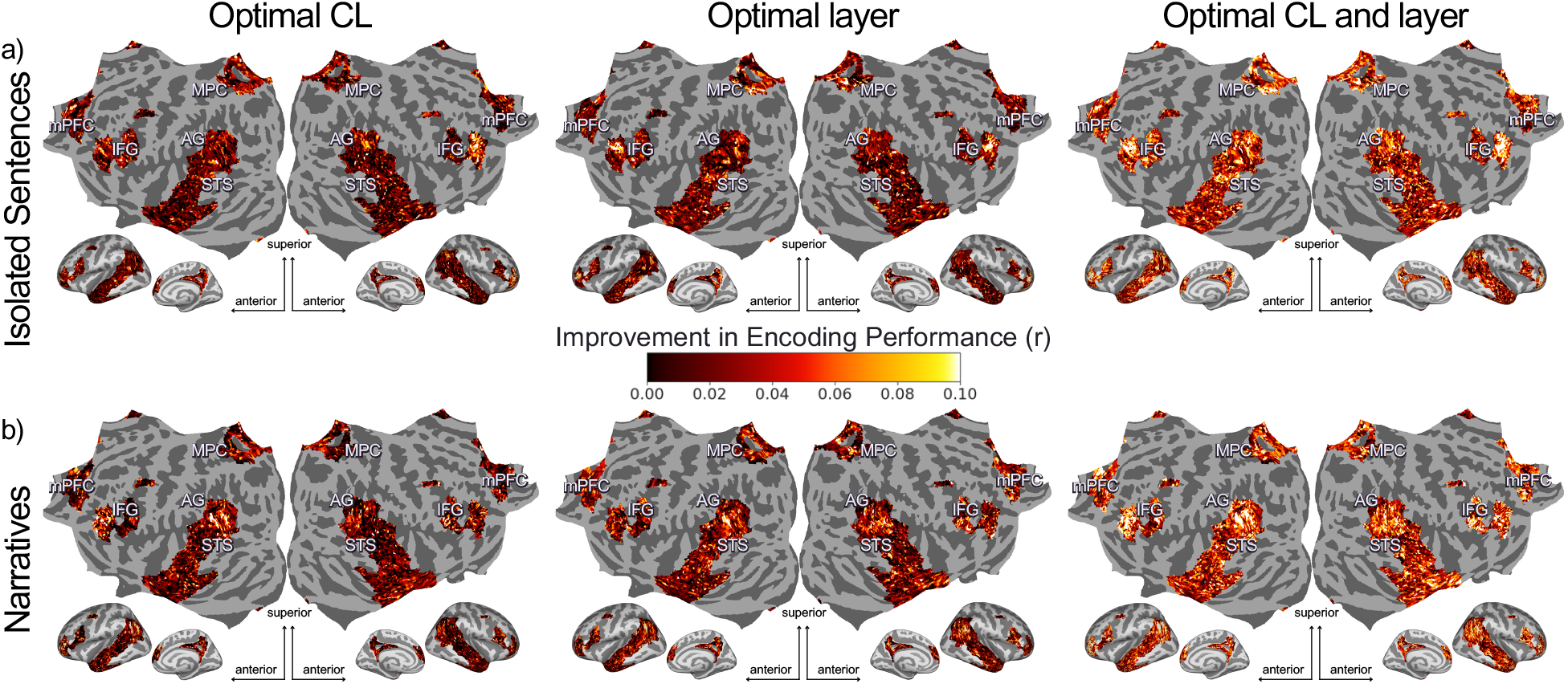
Improvement in encoding performance across language-selective brain regions due to contextual feature optimization. Improvements in brain encoding performance within language-selective regions on a flattened FsAverage cortical surface are shown. Improvement is defined as the difference in encoding performance (r; see Methods) between voxelwise encoding models (VEMs; see Methods) using optimized versus fixed contextual embeddings. Each column corresponds to a different optimization setting: optimal context length (CL), optimal layer, and optimization of both CL and layer. Results are shown for two stimulus conditions: (a) isolated sentences and (b) narratives. Each point represents a cortical vertex within language-selective regions, colored by the magnitude of improvement; voxels outside the language network are shown in gray; mPFC (medial prefrontal cortex), MPC (medial parietal cortex), IFG (inferior frontal gyrus), AG (angular gyrus), and STS (superior temporal sulcus). The largest improvements are observed in bilateral angular gyrus, medial parietal cortex, and inferior frontal gyrus, indicating these regions benefit most from voxel-specific optimization of contextual embeddings. All results in this figure use contextual embeddings from the English BERT model, with fixed features defined as context length 20 and layer 8. Results are shown for Participant 1. Results for other participants and the GPT-2 model are provided in the Appendix Section B and C respectively. Additionally, cortical surface maps showing encoding improvement for all voxels are presented in Appendix Section B.

For both isolated sentences and narratives conditions, optimizing either context length or model layer individually led to participant-specific patterns of improvement, with no single brain region consistently showing the highest improvement across participants. In contrast, joint optimization of context length and model layer showed improvements in most language-selective regions across all participants. In particular, high gains in contextual encoding performance were consistently observed in bilateral angular gyrus, medial parietal cortex, inferior frontal gyrus, and medial prefrontal cortex. These regions are known to support higher-level language and narrative comprehension processes.

Results for the GPT-2 model are provided in the Appendix Section C.

## Discussion and Conclusion

In this study, we investigate whether optimizing contextual semantic embeddings for individual voxels improves their ability to predict brain activity during language comprehension. We optimize contextual embeddings by optimizing the context length, the model layer, or by jointly optimizing both. We evaluate the improvement in contextual encoding performance across two stimulus conditions: isolated sentences and narratives.

Our analysis reveals that voxel-specific optimization of contextual semantic embeddings reliably improves contextual encoding performance across both stimulus conditions. Joint optimization of context length and model layer yields greater improvements than optimizing either alone. Furthermore, the largest gains are consistently observed in brain regions associated with processing higher-level linguistic features. These findings underscore the limitations of using the same contextual semantic embeddings for all voxels and highlight the value of fine-grained optimization for more accurate mappings between language models and brain activity.

To our knowledge, this is the first study to perform voxel-specific optimization of both context length and model layer for contextual semantic embeddings, and to compare these effects across multiple commonly used stimulus conditions (isolated sentences and narratives). This study offers a way to further leverage advances in natural language processing for neuroscience research on semantic representations in the brain.

## Limitations and Future Work

A key limitation of this study is the small dataset (three participants). This is due to the unique nature of the dataset: each participant read text in three stimulus conditions designed from the same set of narratives. For each participant, we had limited training data (30–40 minutes) per stimulus condition, which constrains the observed performance gains. Expanding to a larger dataset would allow for better generalization. Future work could use these optimized contextual embeddings to examine how semantic tuning changes between isolated sentences and narratives.

## Acknowledgment

This work was funded by grants from the German Federal Ministry of Education and Research (BMBF; Grant no. 01GQ1906) and the European Research Council (ERC; Grant no. 101042567).

